# Health-associated gut bacteriocins target TLR4 to suppress intestinal inflammation

**DOI:** 10.64898/2026.07.22.739491

**Authors:** Yuqi Shi, Xiang Fang, Xiaoqian Lin, Xiaoting Xie, Xiaoxuan Chen, Dengwei Zhang, Xin Ma, Jianyu Chen, Xueying Wei, Jiahao Ren, Gengfan Wu, Chang Zhou, Nanzhu Chen, Nan Liu, Guan Yang, Yong-Xin Li

**Affiliations:** Department of Chemistry, The University of Hong Kong, Pokfulam Road, Hong Kong, China; Department of Infectious Diseases and Public Health, City University of Hong Kong, Kowloon, Hong Kong, China; School of Biological Sciences, The University of Hong Kong, Pokfulam Road, Hong Kong, China

## Abstract

The human microbiome maintains host immune homeostasis by secreting bioactive metabolites. However, extending beyond well-characterized metabolites, the functions of most microbiome-encoded peptides remain poorly defined. In this study, we developed a multi-cohort metagenomic framework to profile protective class II bacteriocins—unmodified, ribosomally synthesized peptides—that are enriched in healthy individuals but depleted in patients with inflammatory bowel disease (IBD). We have designated these health-associated bacteriocins as gutcins. Two gutcins, which lack canonical antimicrobial activity, potently attenuate intestinal inflammation in murine models. Cryo-electron microscopy (cryo-EM) reveals that one gutcin, named gutcin 03, directly engages the C-terminus of TLR4, blocking its dimerization and downstream inflammatory signaling. Guided by this structural interface, we generated truncated variants with enhanced potency, demonstrating the amenability of these simple peptides to rational optimization. Collectively, our findings reposition class II bacteriocins from antimicrobial agents to endogenous immunomodulatory effectors and establish a structural and mechanistic foundation for their development as next-generation therapeutics for inflammatory intestinal disorders.

## Introduction

The human gut microbiome profoundly influences host physiology through the production of diverse bioactive metabolites^1^. Deciphering the molecular basis of this host-microbe crosstalk is essential for advancing beyond compositional correlations and realizing the therapeutic potential of the microbiota^2^. Short-chain fatty acids (SCFAs) produced by gut species *Faecalibacterium prausnitzii* and *Akkermansia muciniphila* reinforce the intestinal barrier and promote regulatory T cell differentiation^3–5^, whereas secondary bile acids derived from *Clostridium* clusters modulate host metabolism and immune tone^6,7^. These microbiota-derived metabolites are closely associated with host immune homeostasis. However, upon the onset of disease-associated gut dysbiosis, significant alterations and interaction effects in these metabolites typically occur.

IBD, a chronic intestinal disorder tightly linked to host immune homeostasis and gut dysbiosis^8^. While shifts in microbial composition are well documented in IBD, the functional consequences of these changes, specifically the loss or gain of specific bioactive mediators, are only beginning to emerge as a result of advances in omics technologies^9,10^. Seminal examples include a pro-inflammatory polysaccharide that drives Crohn’s disease flares and indole acrylic acid, a tryptophan metabolite that reinforces epithelial barrier integrity and suppresses inflammation^8,13^. These discoveries underscore the therapeutic promise of microbial metabolites but also expose a critical blind spot: the contribution of healthy gut microbiome-encoded peptides, to intestinal homeostasis and disease remains virtually unknown, largely due to technical challenges in fecal peptide analysis^12^. This gap is especially significant because peptides frequently act as direct effectors of host-microbe interactions and offer favorable features for therapeutic development. Understanding whether and how the gut microbiome peptidome contributes to intestinal homeostasis and disease is therefore a critical priority.

Among gut microbiome-derived peptides, class II bacteriocins are of particular interest. These unmodified, ribosomally synthesized molecules are widely distributed and highly abundant within the human gut microbiota^13^, yet they have historically been considered primarily as antimicrobial agents. However, several observations challenge this limited perspective: certain bacteriocins maintain epithelial integrity^14^ or modulate immune responses^15^, and they share significant structural homology with endogenous host defense peptides^16^. Moreover, their simple biosynthetic logic and the absence of post-translational modifications render them genetically tractable and synthetically accessible, positioning them favorably for drug discovery^17^. Despite these suggestive features, a systematic evaluation of their role in intestinal homeostasis is lacking, and their direct host receptors and downstream signaling pathways remain unknown.

Here, we combine large-scale metagenomic mining with functional and structural analyses to investigate the role of human gut-derived class II bacteriocins. By surveying metagenomes from healthy and diseased cohorts, we identify a set of health-associated bacteriocins that are selectively depleted in IBD. Through bioinformatics-guided chemical synthesis, in vitro screening, and in vivo evaluation, we discovered two anti-inflammatory candidates that act independently of antimicrobial activity to alleviate colitis. Cryo-EM shows that one peptide directly targets the C-terminus of TLR4 to block its dimerization to abrogate inflammatory signaling. Based on this structural interface, we engineer truncated variants with improved activity, revealing that these endogenous peptides provide a tractable scaffold for therapeutic optimization. Our work uncovers a functional class of microbiome-derived immunomodulators and establishes a mechanistic and structural foundation for their translation into novel treatments for inflammatory disorders in the future.

## Results

### Class II bacteriocins are depleted in patients with IBD and suppress macrophage inflammatory responses

To systematically discover human microbiome-derived class II bacteriocins, we constructed a comprehensive reference database by screening 807,469 bacterial genomes, yielding 32,735 non-redundant sequences by IIBacFinder^13^. We then profiled the distribution of these peptides across five human cohorts (four metagenomic and one metatranscriptomic) encompassing 699 healthy individuals and 708 patients with IBD **(Fig. 1a)**, to identify health-associated bacteriocins. Bacteriocin repertoires exhibited significantly higher richness and diversity in healthy individuals compared to IBD patients across all cohorts **(Fig. 1b)**, at both genomic level and in terms of transcriptional activity. We therefore hypothesized that bacteriocins enriched in healthy individuals might contribute to host homeostasis and protection against IBD. To identify such candidates, we compiled 862 bacteriocins that were significantly enriched in healthy individuals. These health-associated peptides were predominantly encoded by beneficial taxa, including *Agathobacter rectalis*, *Roseburia inulinivorans*, and *Anaerobutyricum hallii*, with the production of these bacteriocins emerging as another distinct factor supporting human health and intestinal barrier functions ^18–20^.

**Figure 1.**
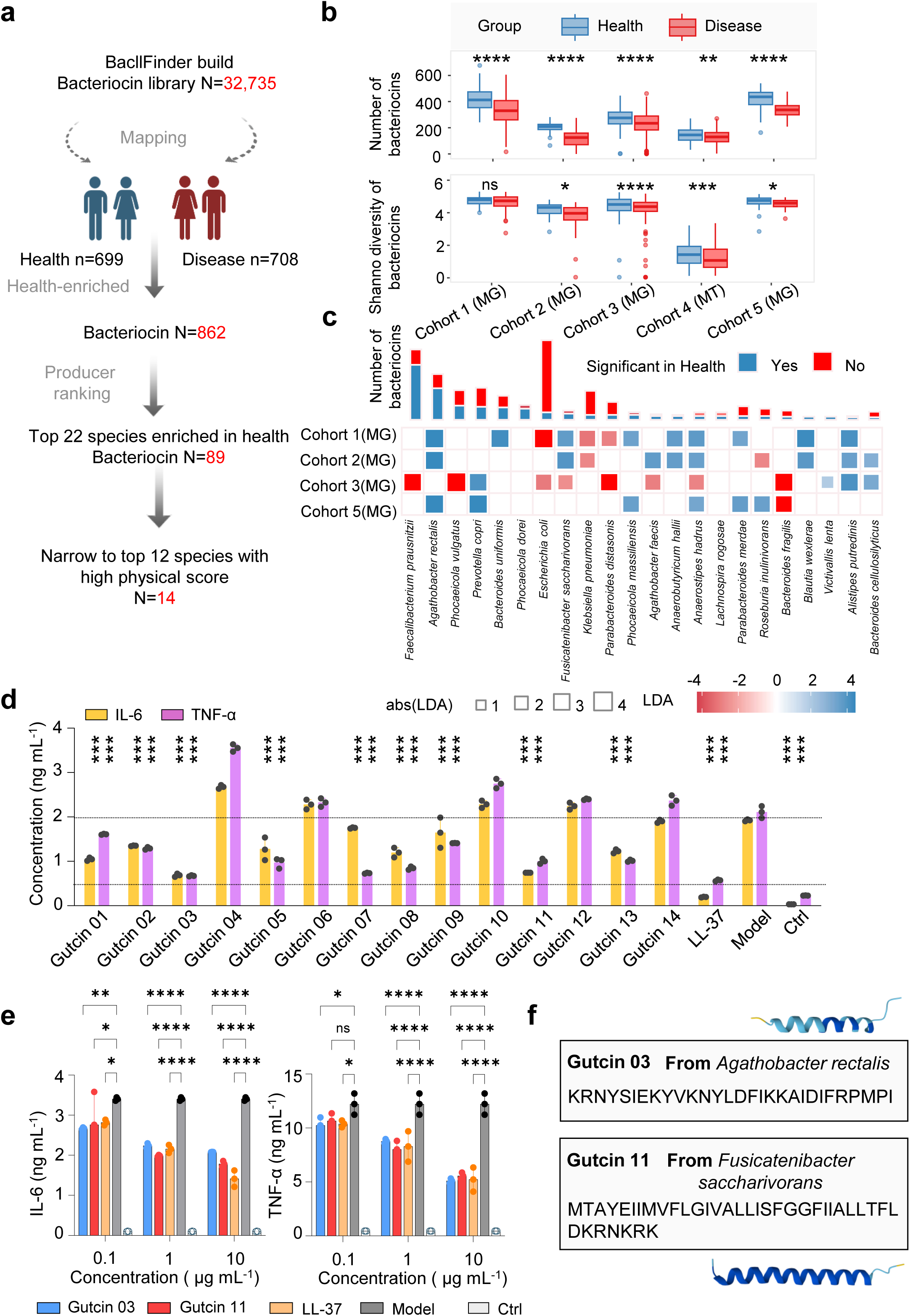
A metagenomics-guided workflow identifies health-associated class II bacteriocins as candidate bioactive molecules. **a,** Schematic overview of the discovery pipeline for class II bacteriocins. **b,** Box plots showing bacteriocin richness and diversity across five independent cohorts. Disease and health groups are represented by red and blue boxes, respectively. **c,** Prioritization of bacteriocin-producing species associated with health. The enrichment status of both the species and their encoded bacteriocins was evaluated across health and disease populations. Bar plots indicate the number of enrichment bacteriocins, whereas the heat map depicts the enrichment profiles of the corresponding bacteriocin-producing species. Blue indicates significant enrichment in health, whereas red indicates significant enrichment in disease. **d,** Primary screening of candidate bacteriocins in BMDMs. Cells were pretreated with individual bacteriocins (10 µg mL^-1^) for 30 min prior to stimulation with LPS (1 µg mL^-1^). Secreted IL-6 (yellow) and TNF- α (purple) levels were quantified by ELISA to assess anti-inflammatory activity. **e,** Dose-dependent anti-inflammatory effects of Gutcin 03 and Gutcin 11. BMDMs were treated with the indicated concentrations (0.1, 1, and 10 µg mL^-1^, of Gutcin 03 (blue), Gutcin 11 (red) and LL-37 (orange; positive control). Cells stimulated with LPS alone served as the model group (dark grey), whereas untreated cells served as the baseline control (light grey). Secreted IL-6 and TNF- α levels were quantified by ELISA to validate the anti-inflammatory activity of the selected bacteriocins. **f,** The producer strains and corresponding peptide sequences of Gutcin 03 and Gutcin 11. For bar graphs, data are shown as mean ± s.e.m. (error bars). For box plots, centre lines indicate the median, box boundaries represent the first and third quartiles, whiskers extend to 1.5× the interquartile range, and points denote outliers. For each treatment condition, n = 3 independent biological replicates. Statistical significance was determined using two-way ANOVA. Significance levels are defined as \**P* < 0.05, \*\**P* < 0.01, \*\*\**P* < 0.001, and \*\*\*\**P* < 0.0001. Source data are provided in the Source Data file.

To further prioritize candidates with potential host-protective functions, we focused on the top 22 producer species consistently enriched across multiple cohorts, narrowing the candidate set to 158 sequences **(Fig. 1c)**. Next, length-based filtering was applied based on the established size ranges of known antimicrobial bacteriocins (20-60 aa) and anti-inflammatory peptides (5-50 aa). Restricting the selection to the overlapping 20-50 aa interval reduced the candidate set to 89 sequences. We subsequently excluded long Sec-dependent variants, retaining only double-glycine and leaderless bacteriocins, which culminated in a final set of 64 sequences^13^. These 64 health-enriched peptides were collectively designated as “gutcins”.

To prioritize candidates for experimental validation, we developed a multi-parameter scoring system that incorporates physicochemical descriptors linked to antimicrobial action (GRAVY hydrophobicity, net charge, terminal hydrophobicity ratio, and amphipathic score) and host-target binding potential (Boman index), as well as oxidizable residue counts and toxicity predictions. This approach ranked candidates from the top twelve health-associated species, yielding 21 high-confidence hits. Guided by biosynthetic structural predictions, we successfully synthesized 14 gutcins for subsequent functional characterization (**Fig. 1a**).

We next investigated whether these peptides modulate host inflammatory responses. Macrophages are a classic cell type in inflammation research, largely due to their high plasticity in polarization and stable inflammatory responses. LPS-stimulated macrophages provide a well-established model for TLR4-driven intestinal inflammation^21,22^, we therefore screened the 14 gutcins for suppression of tumour necrosis factor-α (TNF-α) and interleukin-6 (IL-6) in primary mouse bone marrow-derived macrophages (BMDMs) at 10 µg mL^-1^. In the absence of antibacterial activity, 9 of the 14 gutcins (64%) nevertheless suppressed at least one proinflammatory cytokine, revealing a direct immunomodulatory function **(Fig. 1d)**. Notably, gutcin 03 and gutcin 11 suppressed both TNF-α and IL-6 to an extent comparable to LL-37, a well-characterized host defense peptide with dual antibacterial and immunoregulatory functions^23^. These two candidates were therefore prioritized for detailed characterization. We also evaluated the concentration dependence of this suppression; dose-response experiments revealed that gutcin 03 and gutcin 11 maintained profound inhibition of IL-6 and TNF-α in BMDMs even at concentrations as low as 0.1 µg mL^-1^ **(Fig. 1e)**. Overall, these data demonstrate that gutcin 03 and gutcin 11 modulate macrophage inflammatory responses in a concentration-dependent manner.

### Gutcins alleviate systemic inflammation and ameliorate DSS-induced colitis

To evaluate the in vivo anti-inflammatory efficacy of gutcin 03 and 11, we first established a systemic inflammation model by intraperitoneal (i.p.) challenge of C57BL/6J mice with *E. coli*-derived LPS (5 mg kg^-1^). Mice received i.p. pretreatment with gutcin 03 or gutcin 11 at 50 mg kg^-1^, 2 h prior to LPS administration (**Fig. 2a**). At 6 h post-challenge, histological examination of ileal sections revealed that vehicle-treated mice developed severe intestinal inflammatory lesions, whereas gutcin pretreatment substantially preserved villus architecture and crypt depth (**Fig. 2b, c)**. ELISA quantification confirmed that tissue levels of TNF-α and IL-6 in the ileum and colon were significantly reduced in mice receiving 50 mg kg^-1^ of gutcin (**Fig. 2d, e**). Consistently, serum cytokine levels were significantly lower in gutcin 03 and 11-treated mice across both inflammatory readouts (**Fig. 2f**).

**Figure 2.**
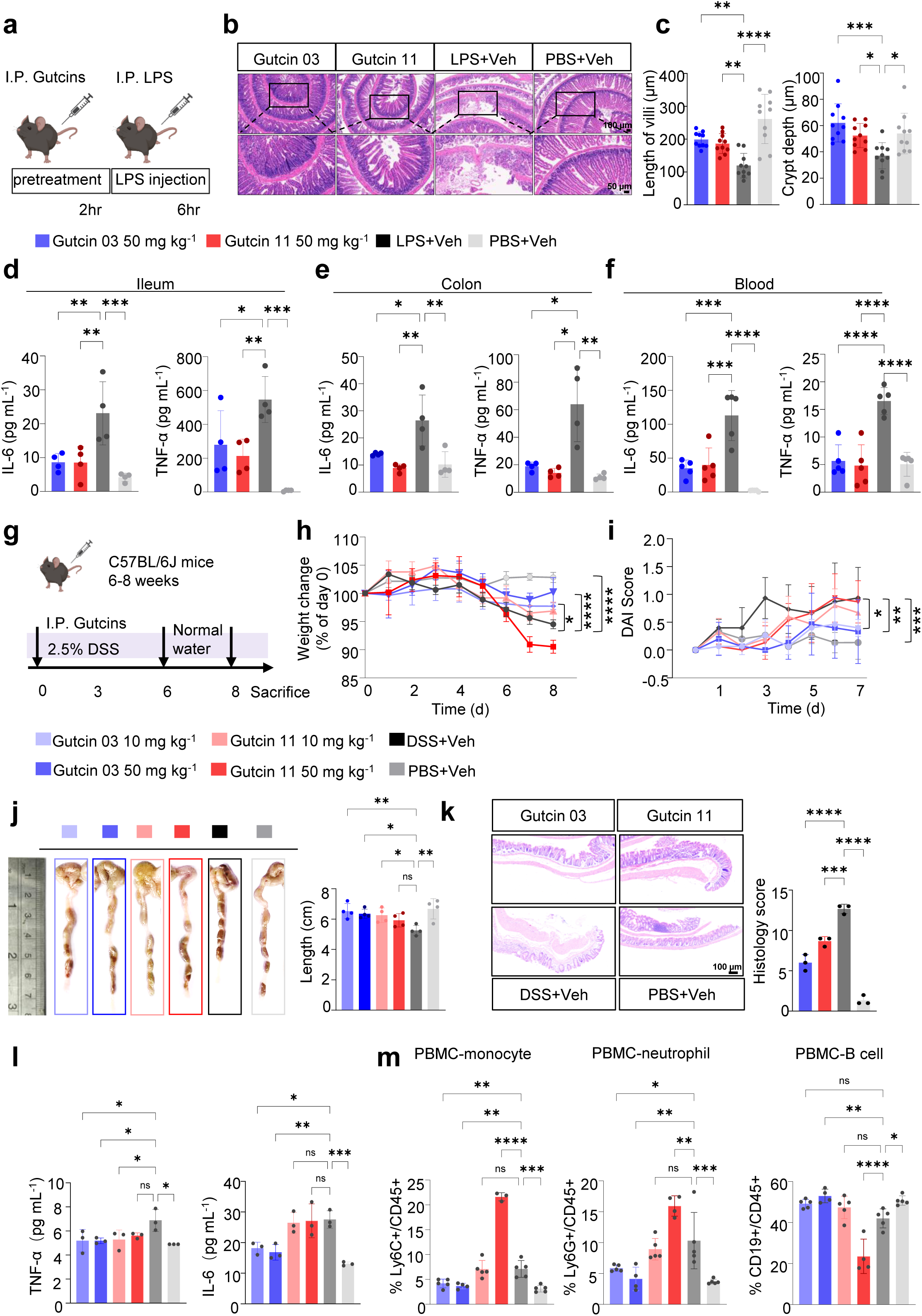
Gutcin 03 and Gutcin 11 suppress systemic inflammation and alleviate experimental colitis in vivo. **a,** Experimental design of the LPS-induced endotoxemia model. C57BL/6J mice received i.p. injections of Gutcin 03 or Gutcin 11 at 50 mg kg^-1^ body weight. Two hours later, systemic inflammation was induced by i.p. administration of *E. coli*-derived LPS (5 mg kg^-1^). Blood and intestinal tissues were collected 6 h after LPS challenge. Each group contained five biologically independent mice, and the experiment was independently repeated three times. **b,** Representative H&E staining of ileal tissues from the indicated groups. Scale bars: 100 µm and 50 µm. **c,** Histopathological alterations were evaluated, and ileal villus length and crypt depth were quantified in ten randomly selected, blinded fields of view using ImageJ (n = 10 measurements per group). **d,** Inflammatory cytokine levels in ileal tissue lysates, **e,** colonic tissue lysates, and **f,** serum measured by ELISA following LPS challenge (n = 4 biologically independent mice per group). **g,** Experimental design of the DSS-induced acute colitis model. Mice received 2.5% (w/v) DSS in drinking water for 6 consecutive days with daily i.p. administration of Gutcin 03 or Gutcin 11 at 10 mg kg^-1^ and 50 mg kg^-1^ body weight. On day 7, DSS was replaced with normal drinking water and gutcin treatment was terminated. Mice were euthanized on day 8 for sample collection and disease assessment. Each group contained five biologically independent mice, and the experiment was independently repeated three times. **h,** Longitudinal changes in body weight during DSS-induced colitis and recovery. Body weight is presented as a percentage of the initial body weight recorded on day 0 (n = 5 biologically independent mice per group). **i,** DAI scores of the indicated treatment groups (n = 5 biologically independent mice per group). **j,** Representative images of colons collected at the experimental endpoint and quantification of colon lengths (n = 4 biologically independent mice per group). **k,** Representative H&E staining of colonic tissues from DSS-treated mice (50 mg kg^-1^). Histopathological evaluation was performed to assess epithelial damage and inflammatory cell infiltration. Scale bars, 100 µm. **l,** Serum concentrations of inflammatory cytokines in DSS-treated mice determined by ELISA (n = 3 biologically independent mice per group). **m,** Flow cytometric analysis of circulating immune cell populations, including monocytes, neutrophils, and B cells, in peripheral blood (n = 5 biologically independent mice per group). For bar graphs, data are shown as mean ± s.e.m. (error bars). Unless otherwise indicated, n represents biologically independent animals. Statistical significance was determined using one-way or two-way ANOVA with appropriate multiple-comparison correction, as detailed in the Methods. Significance levels are defined as \**P* < 0.05, \*\**P* < 0.01, \*\*\**P* < 0.001, and \*\*\*\**P* < 0.0001. Source data are provided in the Source Data file.

We next tested their therapeutic potential in a dextran sodium sulfate (DSS)-induced acute colitis model. Mice received 2.5% DSS in drinking water and daily i.p. injections of gutcin 03 or gutcin 11 at 10 or 50 mg kg^-1^ for 6 days **(Fig. 2g)**. DSS administration caused substantial body weight loss, colon shortening, and severe colonic histopathology. Treatment with gutcins significantly reversed these pathological alterations. Gutcin 03, at both doses, effectively attenuated rapid body weight decline, colon shortening, and disease activity index (DAI) scores during colitis progression (**Fig. 2h-j**). Gutcin 11 exhibited weaker in vivo efficacy than gutcin 03 in acute colitis model. High-dose gutcin 11 (50 mg kg^-1^) failed to restore colon length or prevent weight loss owing to low solubility **(Fig. 2h, j)**.

Histological analysis confirmed reduced muscularis edema and inflammatory cell infiltration in the colons of gutcin-treated mice **(Fig. 2k).** Serum levels of IL-6 and TNF-α were significantly reduced in gutcin 03-treated mice **(Fig. 2l)**. Flow cytometry analysis of peripheral blood immune cells revealed that gutcin 03 (10 and 50 mg kg^-1^) significantly decreased neutrophil and monocyte populations, whereas gutcin 11 at 10 mg kg^-1^ showed a non-significant reduction and the high-dose group paradoxically exhibited proinflammatory effects **(Fig. 2m)**. These data indicate that gutcin 03 possesses superior anti-inflammatory and protective efficacy compared with gutcin 11. Collectively, these findings demonstrate that gutcins 03 and gutcin 11 enriched in healthy individuals was able to alleviate LPS-induced systemic inflammation and gutcins 03 shows protective efficacy in experimental colitis.

To further define the impact of gutcins on local intestinal inflammation and barrier repair, we examined colonic tissues from the DSS model **(Fig. 3a)**. Immunostaining revealed that gutcin treatment substantially reduced colonic IL-6 and TNF-α expression **(Fig. 3b, c)**. Quantitative analysis of Alcian Blue-Periodic Acid-Schiff (AB-PAS) staining showed a significant increase in goblet cell numbers in gutcin 03-treated groups, while gutcin 11 showed an increasing trend compared with DSS-only controls **(Fig. 3d)**. Immunofluorescence analysis further indicated that gutcin 03 and 11 administration markedly increased ZO-1 and Muc2 levels in the colon **(Fig. 3e, f)**. These results indicate that gutcins 03 maintain intestinal homeostasis by attenuating inflammation, preserving the mucus layer, and reinforcing epithelial barrier integrity during colitis, while gutcin 11 exhibits intestinal barrier repair potential.

**Figure 3.**
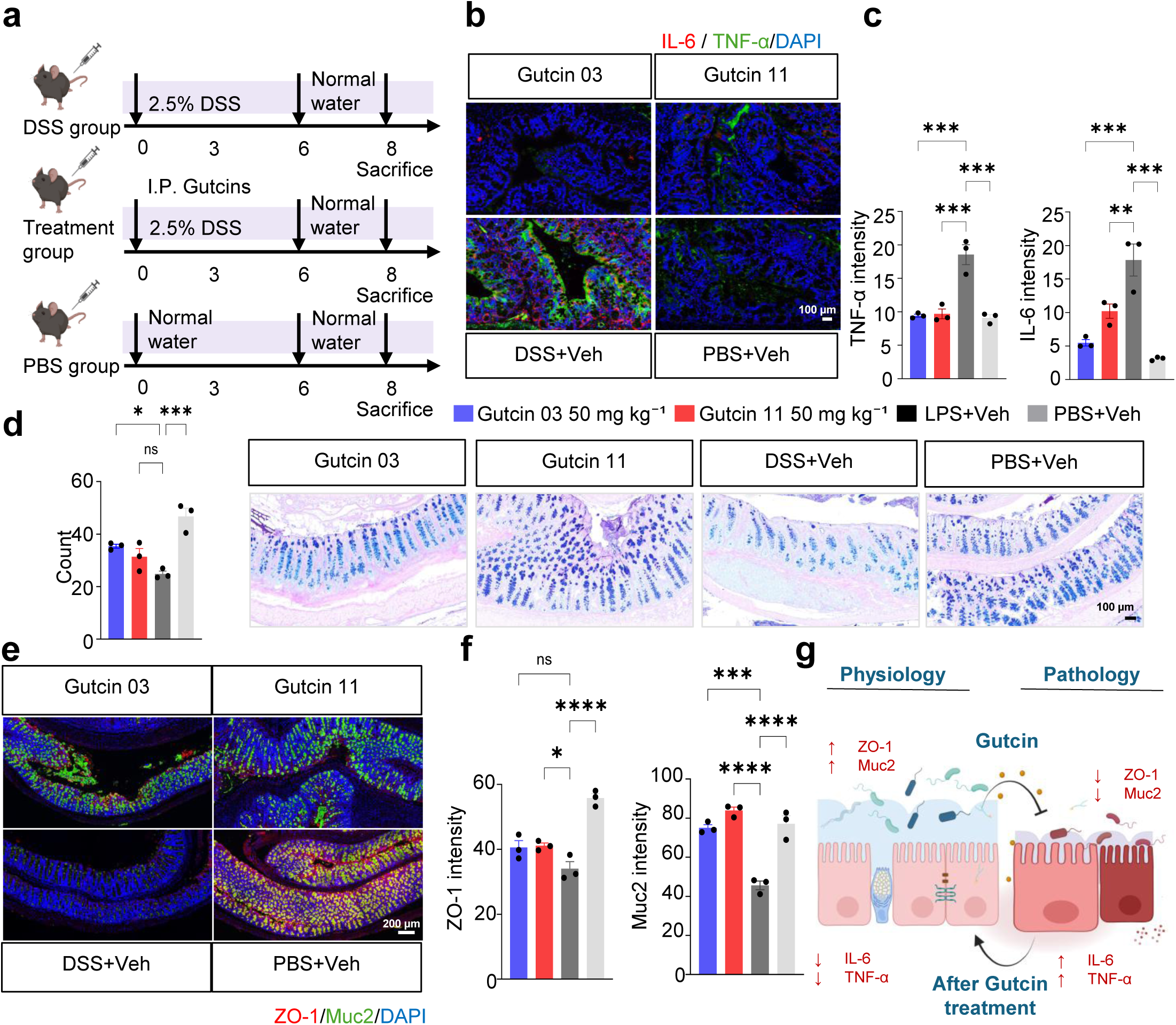
Gutcin promotes the resolution of intestinal inflammation and restores epithelial barrier function. **a,** Experimental design of the DSS-induced acute colitis model. Mice received 2.5% (w/v) DSS in drinking water for 6 consecutive days. During this period, mice were treated daily via i.p. injection with PBS (PBS group), 50 mg kg^-1^ Gutcin 03, or 50 mg kg^-1^ Gutcin 11 (treatment groups), while the DSS-alone group received an equivalent volume of PBS to control for injection stress. On day 7, DSS was replaced with normal drinking water and all treatments were terminated. Mice were euthanized on day 8 for sample collection (n = 5 per group, repeated three times). **b,** Immunofluorescence staining of TNF-α (green) and IL-6 (red) in colonic tissues from DSS-treated mice. Representative images are shown for the PBS group, DSS-alone group, DSS + Gutcin 03 (50 mg kg^-1^), and DSS + Gutcin 11 (50 mg kg^-1^) groups. Scale bars, 100 µm. **c,** Fluorescence intensity of (b) was quantified using ImageJ from randomly selected fields in each group. **d,** Representative photomicrographs of AB-PAS stained colonic sections from the four groups (right), with quantification of goblet cell numbers (left) from randomly selected fields in each group. Goblet cell abundance were assessed histologically in a blinded manner. Scale bars, 100 µm. **e,** Immunofluorescence staining of the epithelial tight junction protein ZO-1 (red) and the goblet cell marker Muc2 (green) in colonic tissues from the four groups: PBS group, DSS-alone, DSS + Gutcin 03 (50 mg kg^-1^), and DSS + Gutcin 11 (50 mg kg^-1^). Representative images from one mouse per group. Scale bars, 200 µm. **f,** Fluorescence intensity of ZO-1 (red) and Muc2 (green) in (f) was quantified using ImageJ from randomly selected fields in each group. **g,** Schematic illustration of Gutcin therapy suppressing intestinal pro-inflammatory cytokines (IL-6, TNF-α) and upregulating Muc2 and tight junction protein ZO-1. These effects collectively accelerate intestinal barrier restoration and mucosal healing. Schematic created with <u>BioRender.com</u>. For bar graphs, data are presented as mean ± SEM. Unless otherwise indicated, n represents biologically independent animals. Statistical significance was determined using one-way or two-way ANOVA with appropriate multiple-comparison correction, as detailed in the Methods. Significance levels are defined as \**P* < 0.05, \*\**P* < 0.01, \*\*\**P* < 0.001, and \*\*\*\**P* < 0.0001. Source data are provided in the Source Data file.

Collectively, these findings demonstrate that gutcins dampen inflammation in both epithelial and lamina propria compartments during colitis, thereby mitigating mucosal damage and promoting barrier repair **(Fig. 3g)**.

### Gutcins suppress TLR4 signaling through direct engagement of the TLR4-TLR4 dimerization interface

Given the established role of aberrant TLR signaling in IBD pathogenesis^24^, we hypothesized that gutcins exert their anti-inflammatory effects by directly modulating TLR pathways. We performed molecular docking against seven candidate receptors, including TLR2 (as monomers and as heterodimers with TLR1 or TLR6), TLR3, TLR4 and TLR7. Docking simulations identified TLR2 (3A7B) as having the highest computed affinity for gutcin 03 (ΔG = -3.12 kcal mol^-1^), followed by TLR4/MD-2 (3FXI; ΔG = -2.5 kcal mol^-1^; **Fig. 4a**). To experimentally validate the target specificity, we treated BMDMs with gutcin 03 in combination with specific TLR agonists. All tested ligands induced robust cytokine production, yet gutcin 03 potently suppressed LPS (TLR4 agonist)-induced TNF-α secretion while only modestly inhibiting Pam3CSK4 (TLR2 agonist)-induced responses (**Fig. 4b**). These findings indicated that TLR4 signaling, rather than TLR2 signaling, is the primary functional target of gutcins in macrophages.

**Figure 4.**
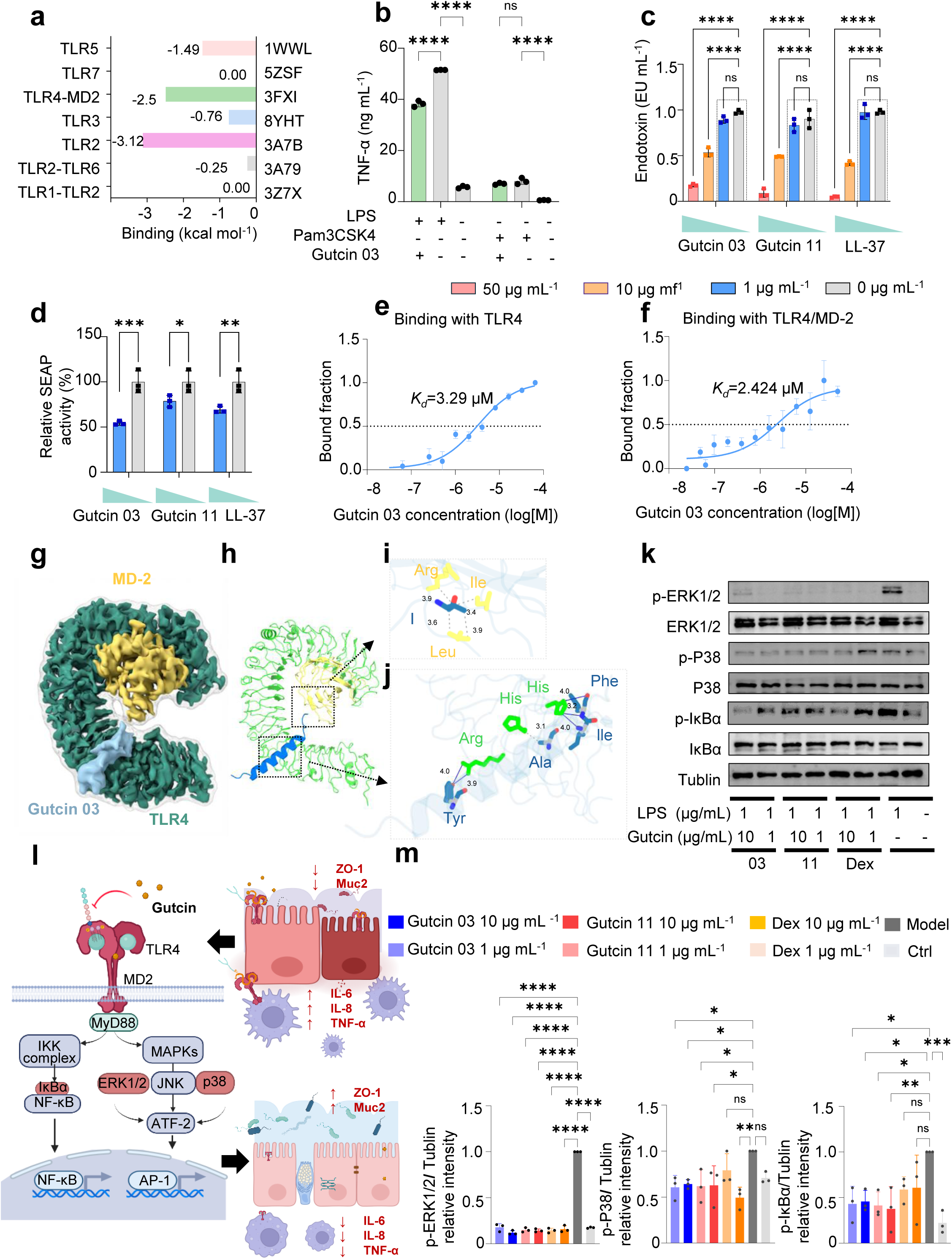
Gutcin promotes the resolution through modulation of Toll-like receptor signalling. **a,** In silico molecular docking analysis of Gutcin 03 with Toll-like receptors (TLRs). Docking scores were calculated using Boltz-2 and Rosetta to evaluate the binding affinity and structural compatibility between Gutcin 03 and individual TLRs. **b,** BMDMs derived from C57BL/6J mice were stimulated with selective TLR2 and TLR4 agonists to activate the corresponding signalling pathways. The inhibitory effect of Gutcin 03 (10 µg mL^-1^) on TNF-α production was subsequently assessed to determine receptor specificity and validate the mechanistic predictions from transcriptomic and molecular docking analyses. **c,** LPS-neutralizing activity of Gutcin 03 and Gutcin 11 at concentrations of 50, 10, and 1 µg mL^-1^, with 0 µg mL^-1^ as a negative control, quantified using the LAL assay. **d,** Inhibition of NF-κB activation in THP-1 NF-κB reporter cells treated with Gutcin 03 or Gutcin 11 at 10 and 1 µg mL^-1^. **e,** Interaction between Gutcin 03 and TLR4 measured by MST. Each experiment was independently performed three times, and Kd values were calculated from globally fitted binding curves derived from all biological replicates. **f,** Interaction between Gutcin 03 and TLR4/MD-2 measured by MST. Each experiment was independently performed three times, and Kd values were calculated from globally fitted binding curves derived from all biological replicates. **g,** Cryo-electron microscopy density map-based structure of Gutcin 03 in complex with the TLR4/MD-2. **h,** Cryo-electron microscopy structure of Gutcin 03 in complex with TLR4/MD-2. **i,** Analysis of the interaction and key residues between MD-2 and Gutcin 03. **j,** Analysis of the interaction and key residues between TLR4 and Gutcin 03. **k,** Immunoblot analysis of IκBα, total and phosphorylated ERK1/2, and total and phosphorylated p38 in cells stimulated with LPS (1 µg mL^-1^) in the presence or absence of Gutcin 03 or Gutcin 11 at the indicated concentrations. Tubulin served as the loading control. **l,** Schematic illustration of the proposed mechanism by which gutcins competitively bind to LPS and the TLR4/MD-2 complex to block TLR4 signalling. This prevents downstream activation of IκBα, ERK1/2, and p38 (highlighted). Consequently, gutcins promote macrophage M2 polarization and Treg differentiation, which collectively account for the observed reduction in intestinal inflammation and the enhanced barrier repair. Adapted from Chen, H. (2024), BioRender.com/v71h443. **m,** Immunoblot analysis was quantified using ImageJ (n = 3 biologically independent experiments). Unless otherwise indicated, data are presented as mean ± s.e.m., and n represents biologically independent samples or animals. Statistical significance was determined using one-way ANOVA or two-way ANOVA. Significance levels are defined as ns, *P* > 0.05; \**P* < 0.05; \*\**P* < 0.01; \*\*\**P* < 0.001; and \*\*\*\**P* < 0.0001. Source data are provided in the Source Data file.

Because TLR4 activation is initiated by LPS binding to MD-2, which triggers receptor dimerization and downstream signaling, we next investigated whether gutcins act by neutralizing LPS. The LAL assay demonstrated LPS neutralization at 50 µg mL^-1^ and 10 µg mL^-1^. However, this neutralizing capacity was nearly abolished at 1 µg mL^-1^ **(Fig. 4c)**. In contrast, an NF-κB reporter assay revealed that both gutcin 03 and gutcin 11 still significantly inhibited NF-κB transcriptional activity even at 1 µg mL^-1^ **(Fig. 4d)**, consistent with our previous observations of cytokine suppression at this concentration **(Fig. 1e)**. These results suggest that LPS neutralization alone cannot account for the anti-inflammatory activity of gutcins at lower concentrations, pointing instead to direct engagement of cellular receptors. To quantify gutcin-TLR4 binding, we performed microscale thermophoresis (MST), gutcin 03 bound to TLR4 and TLR4/MD-2 with dissociation constants (*K_d_*) of 3.29 µM and 2.42 µM **(Fig. 4e, f)**.

To define the structural basis of this interaction, we employed cryo-EM. Gutcin 11 proved refractory to structure determination owing to hydrophobicity-driven self-assembly and poor solubility—consistent with its weaker bioactivity. By contrast, gutcin 03 yielded a high-resolution (3.18 Å) structure of the gutcin 03-TLR4/MD-2 complex **(Fig. 4g, h)**. The density map positioned gutcin 03 within the central region of the TLR4 C-terminal domain, confirming a direct physical association **(Fig. 4g)**. Strikingly, except for the hydrophobic interaction at Ile29, Ile62, Leu69 and Arg72 of MD-2 **(Fig. 4i)**, gutcin 03 forms a hydrogen bond with His429 and hydrophobic interactions with Arg433 and His431 of TLR4, this binding site corresponds to the critical interface that bridges two TLR4 monomers during receptor dimerization **(Fig. 4j)**^25^. This binding mode is mechanistically distinct from previously characterized small-molecule inhibitors, which typically occupy the hydrophobic pocket of MD-2^26^. To our knowledge, this is the first structural determination of a peptide complexed with TLR4/MD-2 and reveals a previously unrecognized inhibitory mechanism that directly blocks the essential TLR4-TLR4 dimerization interface^27,28^.

To validate the functional consequences of this structural blockade, we examined downstream signaling in macrophages treated with gutcins (1 and 10 µg mL^-1^). Both gutcin 03 and gutcin 11 reduced LPS-induced phosphorylation of p38 and ERK1/2 in a dose-dependent manner, with near-complete inhibition at 10 µg mL^-1,^ and also inhibited IκBα degradation **(Fig. 4k, m)**. Together, these data indicate that gutcins block LPS-induced TLR4 activation and downstream kinase phosphorylation through direct engagement of the TLR4 dimerization interface, establishing a structural and mechanistic framework for their anti-inflammatory activity **(Fig. 4l)**.

### Identification of key fragments required for interaction between gutcins and TLR4/MD-2

Having established that gutcin 03 directly engages the TLR4/MD-2 complex, we sought to identify minimal peptide fragments that retain high-affinity binding while improving synthetic accessibility, biochemical stability, and biological potency. Guided by the cryo-EM structure of gutcin 03 in complex with TLR4/MD-2, we performed molecular dynamics simulation (MD) for 1 µs for detailed interaction analysis. For gutcin 03, the most persistent contacts were localized to a central helical segment, which interacts with TLR4 via clustered hydrophobic contacts, hydrogen bonds, and salt bridges at a 16-residue core motif: YVKNYLDIKKAIDIF **(Fig. 4j)**. Lacking a cryo-EM structure of the gutcin 11-TLR4/MD-2 complex, we employed AlphaFold3 to model and predict its binding pose, followed by 1 µs all-atom MD simulations. These trajectories demonstrated stable embedding of gutcin 11 within the ISFGGFIIALLTFLDKR motif. Guided by these structural insights, we identified interacting fragments containing key residues that contribute to binding free energy. We employed two truncation strategies: one directly shortening the core motif and another extending it by three to four residues at either terminus to preserve local helical structure, iteratively optimizing until calculated binding affinity was notably reduced. This approach yielded a library of truncated peptides derived from both gutcin 03 and gutcin 11, with eight candidates exhibiting predicted ΔG < 0 kcal mol^-1^ for the TLR4/MD-2 interaction. To experimentally validate targeting capability, we synthesized the eight candidates and assessed binding by MST. Two variants, 03T1 and 11T3, demonstrated high-affinity binding to TLR4 with *K_d_* values of 1.97 µM and 3.55 µM (**Fig. 5a**). Notably, these two truncated variants bound the TLR4/MD-2 complex more strongly than their full-length counterparts, with *K_d_* values of 504.4 nM and 1.327 µM, confirming that truncation enhanced target affinity **(Fig. 5a)**.

**Figure 5.**
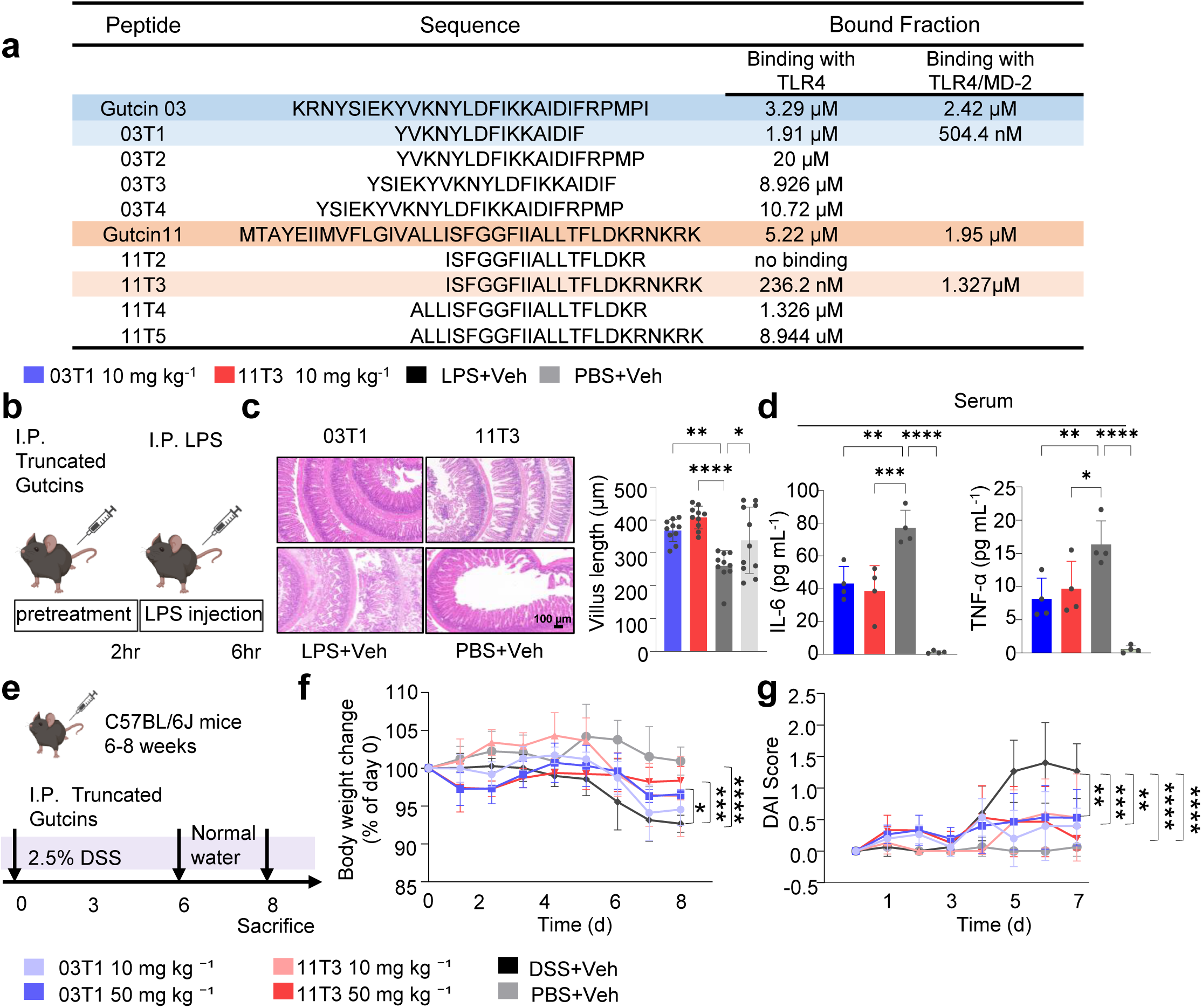
Rationally engineered gutcin-derived fragments retain potent anti-inflammatory efficacy. **a**, Sequences of the truncated gutcin peptides are shown, and their binding affinities to TLR4 and TLR4/MD-2 complex were measured by MST. **b**, Experimental design of the LPS-induced endotoxemia model used to evaluate truncated gutcin derivatives. C57BL/6J mice received i.p. administration of Gutcin 03- or Gutcin 11-derived peptides at 10 mg kg^-1^ body weight. Two hours later, systemic inflammation was induced by i.p. injection of *Escherichia coli*-derived LPS (5 mg kg^-1^). Blood and intestinal tissues were collected 6 h after LPS challenge for histological, immunological, and biochemical analyses. Each treatment group contained five biologically independent mice, and the experiment was independently repeated three times. **c**, Representative H&E-stained sections of ileal tissues from the indicated groups (left), with ileal villus length quantified using ImageJ from randomly selected fields per group (n=10; right). Scale bars, 100 µm. **d**, Concentrations of inflammatory cytokines in serum, quantified by ELISA following LPS challenge (n = 4 biologically independent mice per group). **e**, Experimental design of the DSS-induced acute colitis model. Mice received 2.5% (w/v) DSS in drinking water for six consecutive days together with daily i.p. administration of Gutcin 03- or Gutcin 11-derived peptides at 10 or 50 mg kg⁻¹ body weight. On day 7, DSS-containing water was replaced with regular drinking water and peptide administration was discontinued. Mice were euthanized on day 8 for tissue collection and disease evaluation. Each group contained five biologically independent mice, and the experiment was independently repeated three times. **f**, Longitudinal changes in body weight during DSS-induced colitis and recovery, expressed as a percentage of the initial body weight recorded on day 0 (n = 5 biologically independent mice per group). **g**, DAI scores of the indicated treatment groups (n = 5 biologically independent mice per group). Data are presented as mean ± s.e.m. Unless otherwise indicated, n denotes biologically independent animals. Statistical significance was assessed using one-way or two-way ANOVA with appropriate multiple-comparison corrections, as described in the Methods. Significance levels are defined as \**P* < 0.05, \*\**P* < 0.01, \*\*\**P* < 0.001, and \*\*\*\**P* < 0.0001. Source data are provided in the Source Data file.

### Rationally truncated gutcin candidates function as anti-inflammatory agents in vivo

Encouraged by the enhanced binding affinity of the truncated variants, we next evaluated their anti-inflammatory efficacy in vivo using both LPS-induced systemic inflammation and DSS-induced colitis models. In the systemic inflammation model, 03T1 and 11T3 were administered intraperitoneally at 10 mg kg^-1^ prior to the LPS challenge (Fig. 5b). Histological evaluation of the ileum revealed that vehicle-treated mice developed severe intestinal inflammatory lesions. In contrast, mice pretreated with truncated gutcins at doses of 10 mg kg^-1^, displayed preserved villus length, indicating intact intestinal morphology **(Fig. 5c)**. ELISA of serum, ileum and colon tissues demonstrated that TNF-α and IL-6 levels were significantly reduced in mice treated with of 03T1 or 11T3 at 6 h post-LPS administration **(Fig. 5d)**. Remarkably, before truncation, the parental full-length gutcins exhibited comparable anti-inflammatory efficacy only at the high dose of 50 mg kg^-1^ **(Fig. 2d-2f)**. These findings demonstrate that peptide truncation substantially enhances in vivo biological potency.

Acute colitis was further induced in C57BL/6J mice with 2.5% DSS, and treatment groups received daily intraperitoneal injections of 03T1 or 11T3 at 10 and 50 mg kg^-1^ for 6 days **(Fig. 5e)**. Mice treated with either truncated variant exhibited mitigated body weight loss **(Fig. 5f)**. Both 03T1 and 11T3, at both doses, effectively attenuated DAI scores during colitis progression **(Fig. 5g)**.

Taken together, these results show that truncated gutcins robustly preserve therapeutic efficacy against colitis. Moreover, these findings suggest that even if full-length gutcins are proteolytically degraded within the intestinal microenvironment, the resulting truncated fragments remain highly active and biologically meaningful. The enhanced potency and stability of these rationally designed peptides provide a promising foundation for translatable drug development.

### Gutcins shape the gut microbial community during colitis

To evaluate whether gut microbiome-derived gutcins reshape the intestinal microbiome ecosystem during colitis, we conducted integrated metagenomic analyses on fecal samples from DSS-treated mice **(Fig. 6a)**. Our results reveled that DSS challenge induces a profound collapse in microbial richness and evenness, characterized by a significant reduction in *α*-diversity indices **(Fig. 6b).** Notably, administration of both parent gutcins significantly attenuated this dysbiosis. Metagenomic profiling showed a robust recovery in Chao1 and ACE indices. These data indicate that gutcin 03 and 11-mediated intervention effectively prevents the depletion of microbial diversity, a phenomenon typically associated with acute intestinal inflammation.

**Fig. 6.**
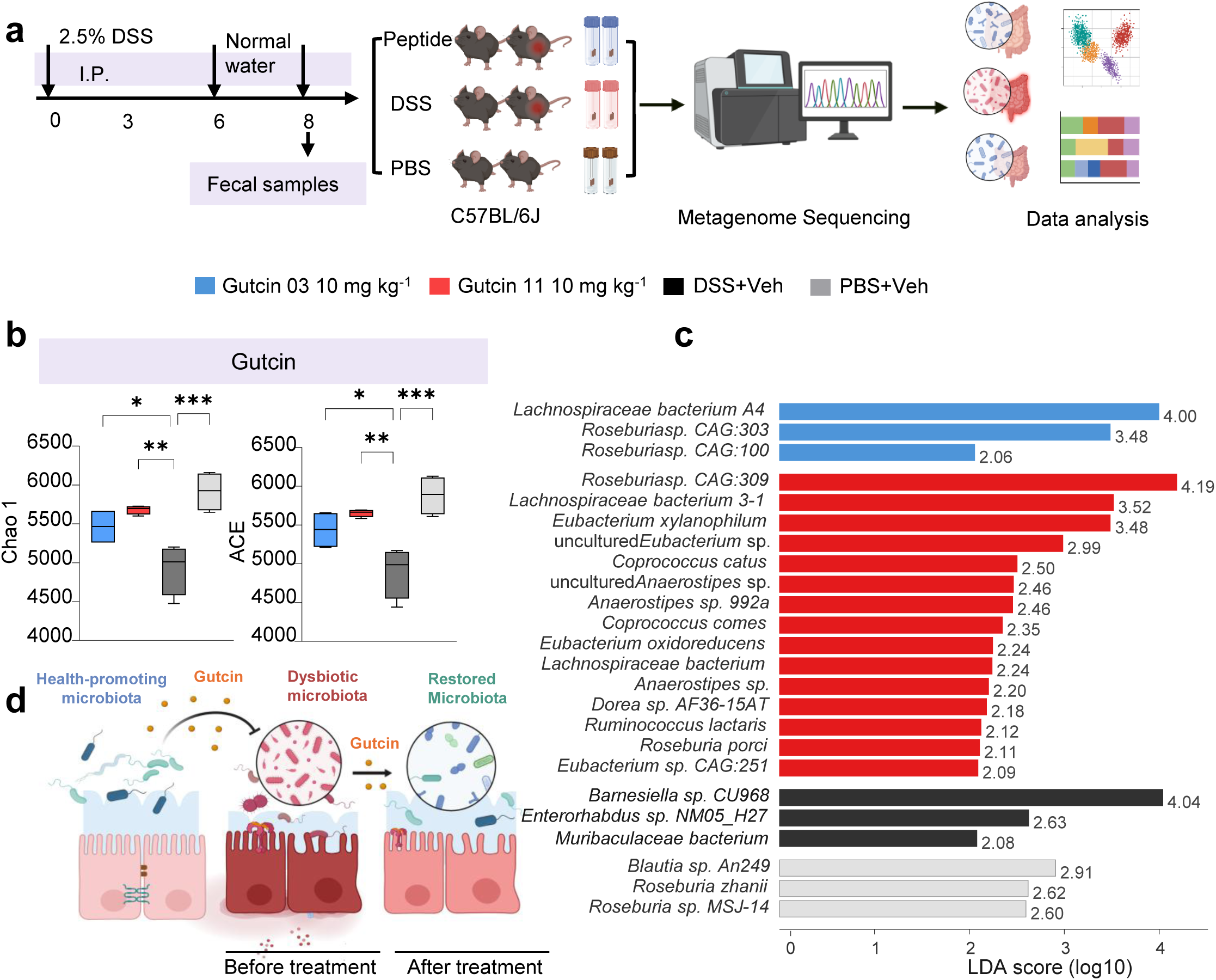
Gutcins and gutcin-derived peptides remodel the gut microbiota and restore microbial homeostasis in DSS-induced colitis. **a,** Experimental design of the DSS-induced acute colitis model used to evaluate the effects of Gutcin 03, Gutcin 11, and their truncated derivatives (03T1 and 11T3; 10 mg kg⁻¹) on gut microbial composition. Fecal samples were collected at the experimental endpoint for shotgun metagenomic sequencing. **b,** Alpha diversity of the gut microbiota in the PBS, DSS, Gutcin 03, and Gutcin 11 groups, assessed using the Chao1 and ACE indices (n = 4 biologically independent mice per group). **c,** LEfSe analysis showing differentially enriched gut bacterial taxa in the PBS, DSS, Gutcin 03, and Gutcin 11 groups. For the Gutcin 03 and Gutcin 11 groups, only beneficial or commensal gut-associated taxa with LDA scores >2 were retained, with a maximum of 15 taxa shown per group. Bar length indicates the LDA score. **d,** Gtcin derived from healthy microbiota effectively repairs gut dysbiosis and facilitates the restoration of disrupted microbiota toward a healthy state. Illustrations created in BioRender: For microbiome analyses, n = 4 biologically independent mice per group unless otherwise indicated. For box plots, centre lines indicate the median, box boundaries represent the first and third quartiles, whiskers extend to 1.5× the interquartile range. Beta diversity was assessed using Bray-Curtis dissimilarity and visualized by PCA. Statistical significance for alpha-diversity indices was determined by one-way ANOVA followed by Tukey’s multiple-comparison test. Differentially abundant taxa were identified using LEfSe. Significance levels are defined as \**P* < 0.05, \*\**P* < 0.01, \*\*\**P* < 0.001, and \*\*\*\**P* < 0.0001. Source data are provided in the Source Data file.

Beyond diversity recovery, LEfSe analysis revealed distinct microbial signatures among the groups, gutcin 03 and gutcin 11 enriched beneficial gut commensals, including *Roseburia*, *Eubacterium*, and *Anaerostipes* **(Fig. 6c)**. In contrast, the DSS group was characterized by inflammation-associated taxa, including *Escherichia coli* and *Enterococcus faecalis*. These findings establish that gutcin 03 and gutcin 11 function as critical modulators of intestinal homeostasis, they do not merely suppress host inflammatory pathways but also act as potent ecological stabilizers that reinforce the resilience of the gut microbial community against colitis-induced dysbiosis **(Fig. 6e)**.

## Discussion

Microbial metabolites are critical regulators of host physiology^12^, yet while small molecules such as SCFAs and secondary bile acids are well established as immunomodulators, the functional roles of microbiota-secreted peptides remain largely unexplored^29^. Through genome mining and multi-cohort metagenomic mapping, we identified a large family of gut-derived class II bacteriocins—termed “gutcins”—that are significantly enriched in healthy individuals and depleted in IBD patients. Two representative candidates, gutcin 03 from *A. rectalis* and gutcin 11 from *F. saccharivorans*, potently attenuated intestinal inflammation in LPS-challenge and DSS-induced colitis models. Class II bacteriocins have been viewed almost exclusively as narrow-spectrum antibiotics since their discovery nearly a century ago^30^, although emerging evidence suggests broader immunomodulatory functions^31^. Our findings establish that these peptides directly target the host TLR4 pathway rather than neutralizing LPS, and structure-guided truncation at the receptor-binding interface further enhanced their activity. Together, these results reveal the therapeutic potential of the human gut peptidome.

Previous studies of commensal metabolites engaging TLR4 have focused on polysaccharides and lipid-like molecules, leaving a gap in our understanding of peptide-mediated modulation^32,33^. For non-human peptides predicted to interact with the TLR4/MD-2 axis, evidence has largely been limited to docking simulations, binding assays or indirect verification through biological experiments^34^. Here, using cryo-EM, we demonstrate that gutcin 03 directly engages the C-terminus of TLR4, modulating receptor dimerization and downstream inflammatory signaling. This mechanism diverges from the prevailing model in which immunomodulatory peptides insert into the hydrophobic pocket of MD-2, thereby expanding the known paradigms of TLR4-ligand interactions. Our structural data not only illuminate the anti-inflammatory mechanism of gutcins but also provide a fresh perspective on how microbial peptides interface with host receptors in the gastrointestinal environment.

Beyond TLR4 signaling, gutcins mitigate mucosal barrier disruption during colitis progression, extending prior observations of barrier protection afforded by class II bacteriocins^35^. Intestinal inflammation driven by dysbiosis elevates LPS levels, activating TLR4 and triggering proinflammatory cytokine release, which, in turn, downregulates ZO-1 and Muc2 expression and compromises epithelial integrity ^35^. Gutcin treatment neutralizes this cascade, reducing inflammation and preserving the mucosal barrier. Mechanistically, we observed that gutcins reduce the proportion of macrophages in the inflamed colonic lamina propria, shifting polarization from the proinflammatory M1 toward the anti-inflammatory M2 phenotype. This M2 polarization promotes IL-10 and TGF-β secretion, which significantly increased the number of Tregs in the colon. demonstrate that bacteriocins ameliorate intestinal inflammation and enhance barrier integrity by orchestrating macrophage polarization and Treg regulation. These coordinated effects support immune resolution and barrier repair. Whether gutcins act directly on T cells or influence other subsets such as Th17 cells warrants further investigation^35–37^.

Gutcin treatment also modulated the gut microbial community during colitis. Administration restricted the expansion of potentially harmful taxa, such as *E.coli* and unclassified *Enterococcus*, while supporting the persistence of beneficial commensals including *Butyricicoccus*, *Ruminococcus*, and *Coprococcus.* This pattern—observed despite the lack of direct antibacterial activity in our assays—suggests that gutcins shape rather than eradicate the microbiota, fostering an environment favorable to SCFA-producing bacteria that may synergize with gutcins to exert immunomodulatory effects^38,39^. However, disentangling direct peptide-mediated effects from indirect shifts driven by altered mucosal ecology requires further investigation.

From a translational perspective, rational truncation of gutcin 03 and gutcin 11 addressed inherent limitations of natural peptides as therapeutic agents^40^. By retaining the core receptor-interaction sites identified through structural studies while discarding non-essential regions, we achieved an optimal balance between synthetic accessibility, stability, and bioactivity. The resulting truncated variants preserved anti-inflammatory efficacy in vivo and exhibited improved drug-like properties, including enhanced binding affinity and proteolytic resistance. These findings underscore the potential of the human microbial metabolite reservoir as a source of adaptable lead compounds amenable to bioengineering for clinical application^41,42^.

The positive correlation between gutcin biosynthetic potential and healthy human cohorts, coupled with our in vivo validation, suggests that gutcin deficiency may contribute to aberrant immune activation in IBD. Conversely, timely supplementation reversed disease pathology, attenuated inflammation and mucosal damage, and promoted recovery. The capacity of these peptides to undergo truncation and modification further highlights their therapeutic promise. Nevertheless, due to limited host availability and the inherent instability of these peptides, we were unable to extensively fractionate intestinal contents to isolate and validate them in situ. As functional omics and high-resolution mass spectrometry continue to advance, capturing these trace microbial peptides under physiologically relevant conditions will expand our understanding of host-microbiota mutualism and provide a broader repository of natural scaffolds for therapeutic optimization in intestinal disorders.

## Methods

### Metagenomics and metatranscriptomics analysis

The raw metagenomic sequencing reads from four cohorts and the raw metatranscriptomic data from one cohort were acquired from NCBI SRA (Sequence Read Archive) under PRJNA400072 (MG, Cohort 1), PRJNA759642 (MG, Cohort2), PRJNA398089 (MG for Cohort 3, and MT for Cohort 4) and PRJEB27928 (MG, Cohort 5). fastp v0.21.1^43^ with default parameters was used to detect and remove low-quality sequencing reads. The sequencing reads belonging to the human host were detected and discarded by kneaddata (https://github.com/biobakery/kneaddata) using a search against the human reference genome T2T-CHM13v2.0^44^. The high-quality sequencing reads retained were subjected to downstream analysis.

### Bacteriocin precursor profiling

Before profiling bacteriocin precursors in metagenomic and metatranscriptomic data, we constructed a reference dataset for class II bacteriocin precursor sequences. IIBacFinder^13^ identified 645,318 high-confidence precursor sequences in bacterial genomes from eight databases, consisting of the RefSeq database (RefSeq, *n* = 257,997), human microbiome (SGB, *n* = 126,702), the Unified Human Gastrointestinal Genome (UHGG, *n* = 285,835), the Human Reference Gut Microbiome (HRGM, *n* = 29,035), global earth microbiome (GEM, *n* = 49,466), ocean microbiome (Ocean, *n* = 21,592), mouse (MGBC, *n* = 26,640) and ruminant gut microbiome (RGM, *n* = 10,211). To remove duplicates and reduce computational load, we deduplicated these sequences at 95% identity and 95% coverage using CD-HIT^45^, resulting in a reference precursor dataset of 32,735 sequences. Then we profiled the distribution of class II bacteriocins across the five processed cohorts.

### Peptide Synthesis

The predicted core peptides of bacteriocins were chemically synthesized via solid-phase peptide synthesis (SPPS) by GL Biochem (Shanghai, China). Their molecular weights were confirmed by mass spectrometry, and purity (determined by high-performance liquid chromatography) was greater than 95%. The synthesized peptide powders were stored at -80 °C and dissolved in dimethyl sulfoxide (DMSO) to 10 mg mL^-1^ before use.

### Mice

C57BL/6J (RRID: MGI: 2159769) mice were maintained under the Laboratory Animal Research Unit of City University of Hong Kong. Six- to eight-week-old mice of both sexes were used. Food and water were provided *ad libitum* throughout this study. All experimental procedures were approved by the Animal Ethics Committee of the City University of Hong Kong (Approval number AN-STA-A-0742 and AN-STA-00001290) and performed in accordance with guidelines from the Laboratory Animal Research Unit of City University of Hong Kong.

### Isolation and culturing of primary macrophages

BMDMs were isolated from the femurs and tibiae of 6- to 8-week-old mice. Bone marrow cavities were flushed with ice-cold PBS using a 25-gauge needle, and the cell suspension was filtered through a 70-μm strainer. After erythrocyte lysis using ACK buffer, cells were resuspended in complete DMEM (high-glucose, 10% heat-inactivated FBS, 1% penicillin-streptomycin) supplemented with 20 ng mL^-1^ recombinant mouse M-CSF (Thermo). Cells were cultured in non-tissue culture-treated dishes at 37 °C with 5% CO2. On day 3, half of the medium (1 mL) was removed and replaced with 1 mL of fresh complete DMEM without M-CSF. On day 7, mature adherent BMDMs were harvested using ice-cold PBS for downstream assays.

### Cell lines

Caco-2, HCT116 and NCM460 cell lines were purchased from the Cell Bank of the Chinese Academy of Sciences (Shanghai, China). Cells were cultured in DMEM (Thermo) supplemented with 10% (v/v) fetal bovine serum (FBS; Thermo), 100 U ml^-1^ penicillin-streptomycin (Beyotime) at 37 °C in a humidified atmosphere with 5% CO₂. All cell lines were routinely tested for mycoplasma contamination and confirmed to be negative using a standard detection kit (Beyotime). Cells were used for experiments within 1 month after resuscitation.

### RNA extraction, qRT-PCR and RNA-seq

Total RNA was extracted from murine intestinal tissues or cell lines using an RNA kit (Vazyme). Complementary DNA was synthesized (Takara) and analyzed by SYBR qPCR (Thermo) with Gapdh as reference (ΔΔ*C*t). Primers are listed in the Supplementary Table.

For RNA sequencing, libraries were prepared from high-quality RNA (RNA integrity number > 8.0). mRNA was enriched, fragmented, and reverse-transcribed. After end repair and adapter ligation, libraries (400-500 bp) were size-selected, amplified and sequenced on an Illumina NovaSeq 6000 platform (Shanghai Personal Biotechnology). Differential expressions were defined as P < 0.05 with a fold change >1.5 or <0.66. Gene ontology enrichment was analyzed using DAVID.

### Microscale thermophoresis (MST) assays

TLR4 (MCE), MD-2 (Nebula Biotechnology), and the TLR4/MD-2 complex (R&D) were prepared at 100 µM in PBS (pH 7.2-7.4). Each protein was labeled with NT-647-NHS ester (Thermo) at room temperature for 1 h under gentle rotation. Unbound dye was removed via ultrafiltration (10 or 30 kDa MWCO), followed by three wash cycles with PBS. The final concentration and labeling efficiency were determined spectrophotometrically. Labeled proteins were serially diluted in MST buffer (PBS supplemented with 0.05% Tween-20) and analyzed using a Monolith NT.115 instrument.

### Molecular docking simulations

The crystal structure of the TLR4/MD-2 complex was retrieved from the Protein Data Bank (PDB ID: 3FXI). Peptide structures were modeled using AlphaFold3^46^, and their binding modes to the TLR4/MD-2 complex were predicted using Boltz-2^47^. The resulting conformations were evaluated using a consensus scoring approach that integrates Boltz-2 confidence metrics and the Rosetta ref2015^48^ all-atom energy function. The top-ranked models were selected for further structural analysis. Molecular visualizations and interaction analyses were performed using PyMOL (version 2.5.5).

### Western blot analysis

Macrophages were stimulated with 1 µg ml^-1^ LPS in the presence or absence of gutcin (1 or 10 µg ml^-1^) for 30 min. Cells were washed with ice-cold PBS and lysed using strong RIPA lysis buffer (Beyotime) supplemented with protease and phosphatase inhibitor cocktails (MCE and Beyotime, respectively). Protein concentrations were determined using a BCA assay. Lysates were separated by SDS-PAGE using precast gels (Vazyme) and transferred to PVDF membranes (Millipore). Membranes were blocked with 5% non-fat milk in TBST for 1 h at room temperature and incubated overnight at 4 °C with the following primary antibodies: phospho-ERK1/2 (1:1,000, ABclonal), ERK1/2 (1:1,000, ABclonal), phospho-p38 MAPK (1:1,000, ABclonal), p38 MAPK (1:1,000, ABclonal), IκBα (1:1,000, ABclonal) and Tubulin (1:1,000, Cell Signaling Technology). Membranes were then incubated with HRP-conjugated goat anti-rabbit IgG (H+L) (1:5,000, ABclonal) for 1 h at room temperature. Chemiluminescent signals were detected using an ECL kit (Thermo) according to the manufacturer’s instructions.

### Dextran sulfate sodium (DSS)-induced acute colitis mouse model

For induction of acute colitis, mice were provided drinking water containing 2.5% (w/v) DSS (MeilunBio) ad libitum for 6 consecutive days, after which the DSS-containing water was replaced with normal drinking water until the experimental endpoint. Gutcins or their truncated derivatives were administered intraperitoneally once daily during DSS treatment at the indicated doses, whereas control mice received an equivalent volume of PBS. Body weight, stool consistency, and fecal occult blood were monitored daily to calculate the DAI. At the experimental endpoint, colons were collected for colon length measurement, histopathological examination, RNA extraction, or reserved at -80 ℃ for downstream biochemical assays.

### Lipopolysaccharide (LPS)-induced systemic inflammation mouse model

For induction of acute systemic inflammation, mice were intraperitoneally administered gutcins or their truncated derivatives at the indicated doses, whereas control mice received an equivalent volume of PBS. Two hours later, systemic inflammation was induced by intraperitoneal injection of Escherichia coli-derived LPS (Solarbio; 5 mg kg^-1^). Mice were euthanized 5-6 h after LPS challenge, and blood samples were collected for serum preparation. The entire intestinal tract was collected for histopathological examination, RNA extraction, and downstream biochemical analyses. Serum cytokine levels and intestinal inflammatory responses were quantified to evaluate the anti-inflammatory efficacy of gutcins.

### Isolation of PP, MLN and CLP immune cells

Single-cell suspensions from PPs and MLNs were prepared by mechanical disruption through a 70-μm cell strainer. For CLP cells, colons were excised, cleared of luminal contents, opened longitudinally, and sectioned into ∼2 cm fragments. The tissues were incubated in HBSS supplemented with 5% FBS, 10 mM EDTA, and 2 mM dithiothreitol (SEED CHEM) for 15 min at 37°C with agitation to remove the epithelial layer. The remaining tissues were digested in RPMI containing 5% FBS, 2 mg/ml collagenase IV (Thermo) and 50 μg ml^-1^ DNase I (Thermo) for 45 min at 37°C with agitation. Following digestion, cell suspensions were filtered through a 70-μm cell strainer. The cell suspensions were resuspended in 40% Percoll, overlaid onto 80% Percoll, and the cells at the interface were collected after centrifugation at 860 g for 20 mins. Erythrocytes in cell suspensions were lysed using ACK buffer, and the cells were immediately used for downstream experiments.

### Flow cytometry

For staining, dead cells were identified using propidium iodide (Sigma–Aldrich, 537060) or Live/Dead Ghost Dye (Tonbo Biosciences). For surface staining, cells were incubated with fluorochrome-conjugated antibodies against CD45 (BioLegend), CD11b (BioLegend), Ly6G (BioLegend), Ly6C (BioLegend), F4/80 (BioLegend), CD206 (BioLegend), CD86 (BioLegend), CD4 (BioLegend), CD19 (BioLegend), and CD3 (BioLegend) in FACS buffer (PBS containing 2% FBS) for 30 min at 4°C. For intracellular staining, cells were fixed and permeabilized using the Cytofix/Cytoperm Fixation/Permeabilization Kit (BD Biosciences) according to the manufacturer’s instructions and subsequently stained with anti-Foxp3 (eBioscience) antibody in permeabilization buffer for 30 mins at 4°C. Data were acquired using a BD FACSCelesta flow cytometer (BD Biosciences) and analyzed using FlowJo software (version 10.2).

### Histological analysis

Intestinal tissue samples were fixed in 4% formalin, embedded in paraffin and sectioned for haematoxylin and eosin (H&E) staining (Servicebio). Goblet cells were characterized by staining intestinal sections with AB/PAS (Servicebio). Brightfield images were acquired using a NanoZoomer 2.0-HT slide scanner. To quantify goblet cell density, the number of AB/PAS-positive cells was counted and normalized to the length of the villus or crypt area using ImageJ software (v.1.53). All quantitative analyses were performed in a blinded manner on at least three representative images per sample.

### Immunofluorescence

Paraffin-embedded intestinal sections were deparaffinized, rehydrated, and subjected to antigen retrieval in 10 mM citrate buffer (pH 6.0). Tissue sections and 4% PFA-fixed cells were blocked with 5% bovine serum albumin (BSA) containing 0.1% Triton X-100 for 1 h. Samples were incubated overnight at 4 °C with primary antibodies against ZO-1 (Servicebio), IL-6 (Servicebio), TNF-α (Servicebio) and Muc2 (Servicebio) (all 1:500 dilution). After washing, samples were incubated with appropriate ABflo 647- or 488-conjugated goat anti-rabbit IgG secondary antibodies (1:400 dilution, ABclonal). Nuclei were counterstained with DAPI. Images were captured using a Zeiss LSM 900 confocal microscope. For quantitative analysis, three random fields of view were selected from each section and blinded, and fluorescence intensity was quantified using ImageJ software.

### Measurement of inflammatory cytokines

The levels of inflammatory cytokines (TNF-α and IL-6) were quantified using ELISA kits (Thermo) according to the manufacturer’s instructions. For blood samples, serum was collected by centrifugation at 5,000*g* for 15 min at 4 °C and analyzed. For intestinal tissue analysis, samples were homogenized in ice-cold PBS supplemented with a protease inhibitor cocktail (MCE), and supernatants were collected after centrifugation at 9,000*g* for 15 min at 4 °C; cytokine levels were normalized to the total protein concentration (determined by BCA assay). For BMDMs, cytokine levels in culture supernatants were measured directly after stimulation, without further normalization.

### Metagenomic and 16S rRNA sequencing and analysis

The sequencing was performed by Novogene Bioinformatics Institute. Total DNA was extracted from fecal samples using the TIANamp Stool DNA Kit (QIAGEN). For 16S rRNA gene profiling, the V4 region was amplified with primers 515F and 806R and sequenced on the DNBSEQ-G99 platform (BGI) at Novogene Bioinformatics Technology Co., Ltd. (Beijing, China). For the Effective Tags, denoising was performed using the DADA2 module in QIIME2 to obtain initial ASVs (Amplicon Sequence Variants). For metagenomic sequencing, libraries from sheared DNA (∼350 bp) were prepared using the Rapid Plus DNA Lib Prep Kit for Illumina (RK20208,. After library quality control, different libraries were pooled based on the effective concentration and targeted data amount. The 5’-end of each library was phosphorylated and cyclized. Subsequently, loop amplification was performed to generate DNA nanoballs. These DNA nanoballs were finally loaded into a flow cell with DNBSEQ-T7 for sequencing at Novogene Bioinformatics Technology Co., Ltd. (Beijing, China).

Microbial community structure was assessed by principal coordinates analysis (PCoA) based on Bray-Curti’s dissimilarity. Group differences were tested using the anosim function in the vegan R package. PICRUSt2 functional predictions were compared via Mann-Whitney U-tests.

### THP-1 reporter cells assay

THP-1-NF-κB-SEAP reporter cells (InvivoGen) were used in this assay. Peptides at various concentrations were added to 96-well plates, with LL-37 serving as the positive control. Subsequently, 1 × 10⁵ cells were seeded per well and incubated with the peptides or vehicle control (DMSO) for 6 h. LPS (0.1 μg mL^-1^) was then added to stimulate the cells for another 24 h. NF-κB activation was quantified by measuring SEAP activity in the culture supernatant using QUANTI-Blue™ solution. Briefly, 180 μL of QUANTI-Blue™ solution mix was prepared in a new 96-well plate. The culture plate was centrifuged at 300 × g for 10 min, and 20 μL of the supernatant from each well was transferred to the corresponding well of the QUANTI-Blue™ plate. Following incubation at 37 °C for 30 min, SEAP levels were determined by measuring absorbance at 640 nm using a SpectraMax iD3 multi-mode microplate reader.

### Cryo-EM sample preparation

The TLR4/MD-2 (R&D) complex was incubated with gutcin 03 at a 1:10 molar ratio on ice for 1 h. Following incubation, the mixture was applied to homemade graphene-coated grids^49^ and then transferred into a Vitrobot Mark IV (Thermo Fisher Scientific). The grids were blotted for 3 s with a blot force of 1 under 100% humidity at 4 °C, and plunge-frozen into liquid ethane for cryo-EM specimen preparation.

### Cryo-EM data collection and processing

Cryo-EM datasets were collected using a Titan Krios Cryo-TEM (Thermo Fisher Scientific) operating at 300 kV. The microscope was equipped with a Falcon 4i direct electron detector and a Selectris post-column energy filter operating with a 20-eV slit width. A total of 10,570 raw movies were collected and subjected to patch motion correction and patch contrast transfer function (CTF) estimation in cryoSPARC (version 4.7)^50^. Particles were selected using the Blob Picker, and multiple rounds of 2D classification were performed to remove bad particles and contaminants. The remaining clean particles were used for ab initio reconstruction, and 700,851 particles were finally selected for non-uniform (NU) refinement. To refine the bound peptide density, a local mask focusing on this peptide density was generated and imported back into cryoSPARC for focused 3D classification. Finally, 26.2% of the particle set was selected and subjected to a final round of non-uniform refinement and local refinement, yielding a final map with an overall resolution of 3.18 Å based on the gold-standard FSC = 0.143 criterion. To build the model, we first docked the published human TLR4/MD-2 complex structure (PDB ID: 3FXI)^25^ and a short peptide predicted by AlphaFold3^51^ into the 3.18 Å cryo-EM density map using UCSF ChimeraX5^52^ and then refined the resulting model in CryoNet.Refine6.

### Molecular dynamics simulations

Molecular dynamics simulations were performed using the Amber22 package. The structures of TLR4/MD-2 in complex with gutcin 03 were obtained from Cryo-EM structural data, while the structures of gutcin 11 in complex with TLR4/MD-2 were generated using AlphaFold3. The protonation states of titratable residues (His, Glu, Asp) were assigned based on predicted pKa values from the PROPKA80 software, combined with visual inspection of local hydrogen-bonding networks. The AMBER force field ff14SB^53^ was employed for the normal protein residues. Counter ions were added to neutralize the total charge system using the “tleap” module. The resulting system was solvated in a rectangular box of TIP3P^54^ water molecules, extending to a minimum distance of 12 Å from the protein surface. The system underwent energy minimization using steepest descent, followed by conjugate gradient, to resolve steric clashes. Subsequently, the system was gently annealed from 0 to 300 K in the canonical ensemble for 300 ps with a restraint of 100 kcal mol^-1^·Å^-2^ on the protein backbone atoms. For density equilibration under isothermal-isobaric ensemble (NPT) at a target temperature of 300K and a target pressure of 1.0 atm using a Langevin thermostat and a Berendsen barostat. For density equilibration under NPT conditions, a force-constant gradient protocol was implemented across four consecutive 1-ns stages with progressive reduction of harmonic backbone restraints: 100 kcal mol^-1^·Å^-2^ (0-1 ns), 10 (1-2 ns), 1 (2-3 ns), and complete removal (3-4 ns). Finally, 1 μs productive MD simulations under the NPT ensemble were performed. The covalent bonds containing hydrogen atoms were constrained using SHAKE^55^ to enable an integration step of 2 fs. Nonbonded interactions were treated with the Particle Mesh Ewald (PME) method with the cutoff set at 10 Å. The simulations were performed with the GPU version of the AMBER 22 package^56^.

### Statistical Analysis and Reproducibility

Statistical analyses regarding the correlation between biosynthetic enzymes and disease states were performed using publicly available datasets; consequently, parameters such as sample size determination, data exclusion, randomization, and blinding did not apply to this computational re-analysis. For all other experimental assays, sample sizes were determined to ensure adequate statistical power, and statistical outliers were identified and excluded based on Grubbs’ test. For the in vivo animal experiments, mice were randomly assigned to experimental groups and cages, and histopathological scoring was performed in a blinded manner.

## Acknowledgments

This work is partially funded by Hong Kong Research Grants Council Collaborative Research Fund (C7014-24G and C6012-22GF) and General Research Fund (17115322 and 17102123) to Y.X.L.

## Author contributions

Y.L., G.Y., N.L. and Y.S designed the research. Y.S., X.F. and Q.X. performed the screening of gutcins. Y.G., X.F., X.X., X.M. and X.M. conducted experiments with mouse models. Mass spectrometry-based analyses were carried out by N.C. and F.G. Y.S. and X. W conducted the related molecular experiments. Y.S., X.X., C.Z. and X.L. performed computational docking. D.Z. and X.L. performed the metagenomic, RNA-seq and 16S rRNA sequencing data analysis. Y.S., X.F., X.M. and X.X. performed flow cytometry experiments. J.C., R.J. and N.L. performed Cryo-EM data collection. Y.S. prepared all the figures and wrote the paper. All authors analyzed data and discussed results. All authors participated in preparing the manuscript. Y-X.L. supervised the project.

## Additional information

### Supplementary information

#### Code availability

All sequencing data generated in this study have been deposited in NCBI SRA with accession number PRJNA1497627. All other data needed to evaluate the conclusions in the paper are present in the paper and/or the Supplementary Materials.

